# Cyborg islets: implanted flexible electronics reveal principles of human islet electrical maturation

**DOI:** 10.1101/2024.03.18.585551

**Authors:** Qiang Li, Ren Liu, Zuwan Lin, Xinhe Zhang, Israeli Galicia Silva, Samuel D. Pollock, Juan R. Alvarez-Dominguez, Jia Liu

## Abstract

Flexible electronics implanted during tissue formation enable chronic studies of tissue-wide electrophysiology. Here, we integrate tissue-like stretchable electronics during organogenesis of human stem cell-derived pancreatic islets, stably tracing single-cell extracellular spike bursting dynamics over months of functional maturation. Adapting spike sorting methods from neural studies reveals maturation-dependent electrical patterns of α and β-like (SC-α and β) cells, and their stimulus-coupled dynamics. We identified two major electrical states for both SC-α and β cells, distinguished by their glucose threshold for action potential firing. We find that improved hormone stimulation capacity during extended culture reflects increasing numbers of SC-α/β cells in low basal firing states, linked to energy and hormone metabolism gene upregulation. Continuous recording during further maturation by entrainment to daily feeding cycles reveals that circadian islet-level hormone secretion rhythms reflect sustained and coordinate oscillation of cell-level SC-α and β electrical activities. We find that this correlates with cell-cell communication and exocytic network induction, indicating a role for circadian rhythms in coordinating system-level stimulus-coupled responses. Cyborg islets thus reveal principles of electrical maturation that will be useful to build fully functional *in vitro* islets for research and therapeutic applications.

## Main text

Human pluripotent stem-cell-derived islets (SC-islets) hold great promise for advancing diabetes research, pharmacology, and transplantation therapy^1–3^. However, SC-islets often exhibit immature function, marked by poor insulin stimulation capacity^4–6^. To address this issue, various methods have sought to enhance SC-islet functional maturity, including extended culture, cell enrichment, and diurnal feeding-fasting entrainment^7–11^. These approaches trigger postnatal islet maturation hallmarks, typically undone with the loss of glucose responsiveness in diabetics, that improve the ability of transplanted SC-islets to mitigate diabetes. However, *in vitro* SC-islets still exhibit less mature phenotypes than transplanted SC-islets, and both lack the precision, kinetics, and magnitude of hormone secretion of adult islets^12–14^. Whether these limitations reflect poor coordination between (or within) populations of SC-islet cell types, or intrinsic heterogeneity in their maturation, remains unclear. Thus, a system-level, single-cell resolution understanding of how SC-islets turn specialized holds transformational potential to build fully functional SC-islets for research and regenerative medicine applications.

Like neurons, islet α/β cells release cargos (insulin/glucagon) in response to membrane potential changes, which are driven by glucose oxidation-generated ATP^15^. Both the glucose coupling and threshold for hormone release increase during functional maturation^16,17^. Yet, the underlying mechanisms have been challenging to study, as it has been difficult to continuously track islet-wide simultaneous α- and β-cell activities across intact maturing islets. Imaging-based approaches, including Ca^2+^ flux readouts, can only report on one plane of view at a time within the three-dimensional (3D) islet volume^18,19^. Intracellular electrophysiology is incompatible with long term-stable measurements, as it requires cell membrane disruption^15^. Extracellular recordings, as with classic multi-electrode arrays, are non-invasive and continuous, but contact islets only at one surface plane^20,21^. Recently, we synergized advances in soft materials and nanoelectronics to build tissue-like nanoelectronics with subcellular feature size, tissue-level flexibility, and mesh-like networks, enabling seamless integration across 3D tissue^22,23^. Implanting and distributing such nanoelectronics throughout developing organoid bodies, thus forming “cyborg” organoids, allows studying cell-level electrophysiology dynamics during organogenesis^24–27^. These innovations have enabled 3D electrophysiology studies of neurons, but have not been applied to islets, despite their similarities in developmental programming and electrophysiological behaviors^28^.

Here, we develop cyborg SC-islets, enabling continuous single-cell electrophysiology across the intact 3D islet network during functional maturation. Using embedded electrodes, we successfully capture stable extracellular voltage spike dynamics from SC-islets. By adapting spike sorting analytics from neural studies, we attain robust sorting of single-unit action potentials over months of chronic recording. The sorted action potentials discern electrical activities specific to α and β-like (SC-α and β) cells based on their responses to glucose, and further identify two major electrical states for both, marked by distinct glucose thresholds for action potential firing. We trace the evolution and coordination of these electrical states over months of extended culture, when SC-islets develop specialized hormone responses. The population dynamics of SC-α and β glucose-coupled electrical activities reveal widespread heterogeneity. We find that improved glucose responsiveness during this early maturation time course reflects increasing presence of SC-α and β electrical states with low basal firing rates. Through single-cell RNA sequencing (scRNA-seq) of cyborg islets, we further find that these electrophysiological changes are linked to broad induction of energy and hormone metabolism gene pathways across SC-α and β cells. Finally, we show that introducing 24-hour feeding-fasting cycles elicits circadian glucose-coupled insulin/glucagon secretion rhythms via sustained coordination of SC-α and β cell electrical activity oscillations. These changes are underpinned by upregulation of cell-cell communication and exocytic gene networks, revealing a role for circadian rhythms in harmonizing SC-islet-wide α/β-like activities to coordinate stimulus-coupled hormone responses. Together, our findings demonstrate the power of cyborg islets as a unique platform to dissect principles of electrical islet cell maturation, which will inform efforts to build fully mature SC-islets for research and generative medicine applications.

## Results

### Building cyborg human SC-islets

We created cyborg human SC-islets by seamlessly integrating stretchable mesh nanoelectronics with SC-islet cells, enabling chronically stable electrophysiology throughout *in vitro* maturation (Fig. 1a). SC-islet cells are seeded with stretchable mesh nanoelectronics on a Matrigel hydrogel substrate, followed by condensation of the cell-nanoelectronics structure by cell-cell attraction forces. 3D morphogenesis is triggered by co-culture with mesenchymal stem cells (MSCs), which promote a self-organized 3D folding process^24,29^. Electrical recordings were performed weekly over a 2-month time course of early maturation prompted by extended culture, and hourly upon further maturation induced by entrainment to daily feeding cycles (Fig. 1a). The stretchable mesh nanoelectronics were specifically designed^25,27^ to enable localized and cell-level electrophysiology recordings while minimizing noise interference (Extended Data Fig. 1a-c). To evaluate their performance, we assessed electrode impedance (Extended Data Fig. 1d), which showed consistent and stable performance across different samples. Additionally, we conducted long-term experiments in a physiological solution. Our results indicate that the electrodes maintain a stable performance across 10 weeks, ensuring long term-reliable electrical recordings (Extended Data Fig. 1e). These results demonstrate suitability and reliability of the stretchable mesh nanoelectronics for long-term electrical recordings. To generate SC-islet cells, we employed a stepwise 3D differentiation approach involving controlled administration of signaling factors and small molecules in a series of growth media over a 20-day protocol^7,30^. The resulting SC-islets gain glucose-responsive insulin secretion function upon 3-4 weeks of extended culture (in medium without serum or exogenous factors^7–9^. Terminal differentiation-stage SC-islets (see Methods) were then dissociated into single cells and subsequently embedded with 64-channel stretchable mesh nanoelectronics to form cyborg SC-islets (Extended Data Fig. 1f-i).

**Fig. 1.**
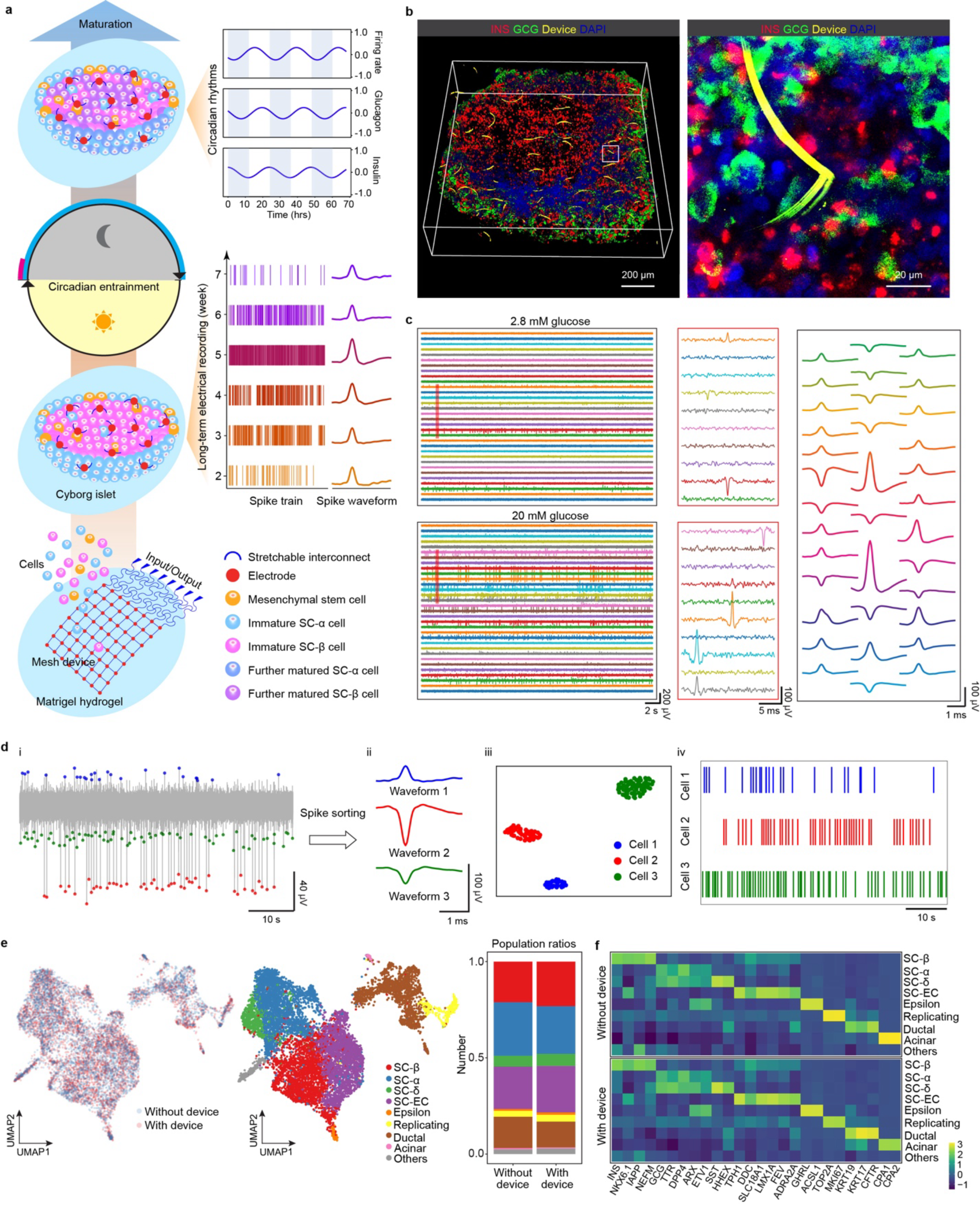
Building cyborg human SC-islets. **a,** Schematics showing integration of stretchable mesh nanoelectronics into cyborg islets for long-term stable electrical recording. Human pluripotent stem cell-derived islet endocrine cells are embedded with stretchable mesh nanoelectronics on a Matrigel hydrogel substrate. Co-culture with mesenchymal stem cells stimulates self-assembly of cyborg islets within 48 hours. Electrical recordings were performed weekly over a 2-month time course of early maturation prompted by extended culture, and hourly upon further maturation induced by entrainment to 24-hour feeding cycles. **b**, Left: Three-dimensional (3D) rendering of fluorescence imaging of a cleared, immunostained cyborg islet at 2 months post device integration. Right: Zoom-in view of the white box inset in the left panel shows intimate coupling of flexible interconnects from the device with SC-α and SC-β cells. INS (Insulin), red; GCG (Glucagon), green; R6G (device), yellow; and DAPI, blue. **c**, Left: Representative raw voltage traces showing spike bursting dynamics for a cyborg islet exposed to 2.8 mM and 20 mM glucose at 6 weeks post device integration. Middle: Single spikes from the regions highlighted in the red box insets from 2.8 mM and 20 mM glucose treatments. Right: Representative averaged single spike waveforms. **d**, Data analysis pipeline for the spike analysis. Spike sorting was performed on 300-3000 Hz filtered voltage traces (i) to identify distinct waveforms (ii). UMAP waveform clustering visualizes cell-specific activity profiles (iii). Representative cell-specific spike trains detected from 1-minute continuous recordings (iv). **e**, 2D UMAP clustering of single-cell RNA expression profiles from cyborg islet (with device) and control islet (without device). Cells are colored by sample of origin (left) and by cell type assignment (middle), with cell type compositions in cyborg and control islets shown to the right. **f**, Expression profiles of markers for each cell type identified in cyborg and control islets, shown as normalized z-scores.

We next sought to conduct immunostaining, electrical recordings, and scRNA-seq to validate the feasibility of cyborg islets. First, immunostaining results demonstrated uniform embedding of the stretchable mesh nanoelectronics throughout the 3D volume of cyborg SC-islets (Fig. 1b, Extended Data Fig. 1l,m). Compared to control SC-islets (Extended Data Fig. 1j,k; without embedded mesh nanoelectronics), cyborg SC-islets show the same composition and distribution of SC-α, -β, and MSC cells (Fig. 1b, Extended Data Fig. 1l). Zoom-in images demonstrate effective coupling of the nanoelectronics with SC-α and SC-β cells (Fig. 1b, Extended Data Fig. 1m), essential for successful cell-level electrical recordings. Second, using the embedded electrodes, we captured voltage spike bursting dynamics in response to low (2.8 mM) and high (20 mM) glucose concentrations, as depicted by representative voltage traces (Fig. 1c, left and middle). The raw electrical traces were subjected to bandpass filtering, limiting the frequencies to the 300-3,000 Hz range. Then, spike sorting was carried out by adapting the MountainSort^31^ and SpikeInterface^32^ algorithms, widely used in neural signal analysis. Specifically, single spike waveforms exceeding the detection threshold were identified (Fig. 1c, right) and clustered. Example of the clusters is visualized using Uniform Manifold Approximation and Projection (UMAP)^33^ (Fig. 1di-iii). We then extracted cell-specific activity profiles from the continuous electrical recordings (Fig. 1div). This methodology enabled us to continuously monitor single-cell electrical activities within SC-islets, offering valuable insights into their maturation dynamics.

As a third validation of the feasibility of our approach, we conducted scRNA-seq on both cyborg and control SC-islets, and investigated the impact of the embedded mesh electronics on cell-type composition and gene expression. Unsupervised UMAP^33^ clustering of gene expression profiles (Fig. 1e) indicated that cells from cyborg and control SC-islets grouped together across all cell clusters. Using previously established gene markers^7,14^, we annotated endocrine and non-endocrine cells as the two major cell clusters, with the same cell types within each cluster for cyborg and control SC-islets (Fig. 1e, f, Extended Data Fig. 2). Endocrine cells included SC-α (*GCG^+^ ARX^+^*), SC-β (*INS^+^ NKX6-1^+^*), SC-EC (*TPH1^+^ FEV^+)^*, SC-δ (*SST^+^ HHEX^+^*), and a small population of SC-ε cells (*GHRL^+^ ACSL1^+^*). A continuous spectrum was observed between SC-β and SC-EC cells, and between SC-α and SC-δ, consistent with emergence from common precursors^34,35^. Non-endocrine cells comprised ductal (*KRT17^+^ KRT19^+^*), acinar (*CPA1^+^ CPA2^+^),* and a replicating population (*TOP2A^+^ MKI67^+^)*. Importantly, we observed highly consistent gene expression patterns between cyborg and control SC-islets across all endocrine and non-endocrine cell types, indicating a negligible effect of nanoelectronics implantation on the SC-islet landscape. Collectively, these results demonstrate the reliability and robustness of the cyborg SC-islet platform for studying SC-islet maturation dynamics *in vitro*.

### Capturing cell type-specific electrical dynamics

Using cyborg islets, we can distinguish islet-wide SC-α and β cellular electrophysiology features (Fig. 2a). β cells are characterized by hyperpolarization and electrical inactivity under physiologically low glucose levels. When the glucose concentration rises, however, ATP generated from glucose oxidation leads to K_ATP_ channel closure, causing membrane depolarization and the initiation of electrical activity that triggers insulin exocytosis^15^. α cells, conversely, fire action potentials under low glucose levels, reflecting activation of voltage-gated Na^+^ and Ca^2+^ and inhibition of K_ATP_ channels^36^. SC-islets also exhibit voltage-gated glucagon and insulin secretion from SC-α and SC-β cells, respectively, in response to low and high glucose concentrations^8,37^. To identify SC-α and β-specific activities, we analyzed electrical activity features recorded from cyborg SC-islet cells, including spike firing rate, amplitude, duration, peak-trough ratio, width at half-maximum, repolarization slope, and recovery slope (Fig. 2b, Extended Data Fig. 3a, b).

**Fig. 2.**
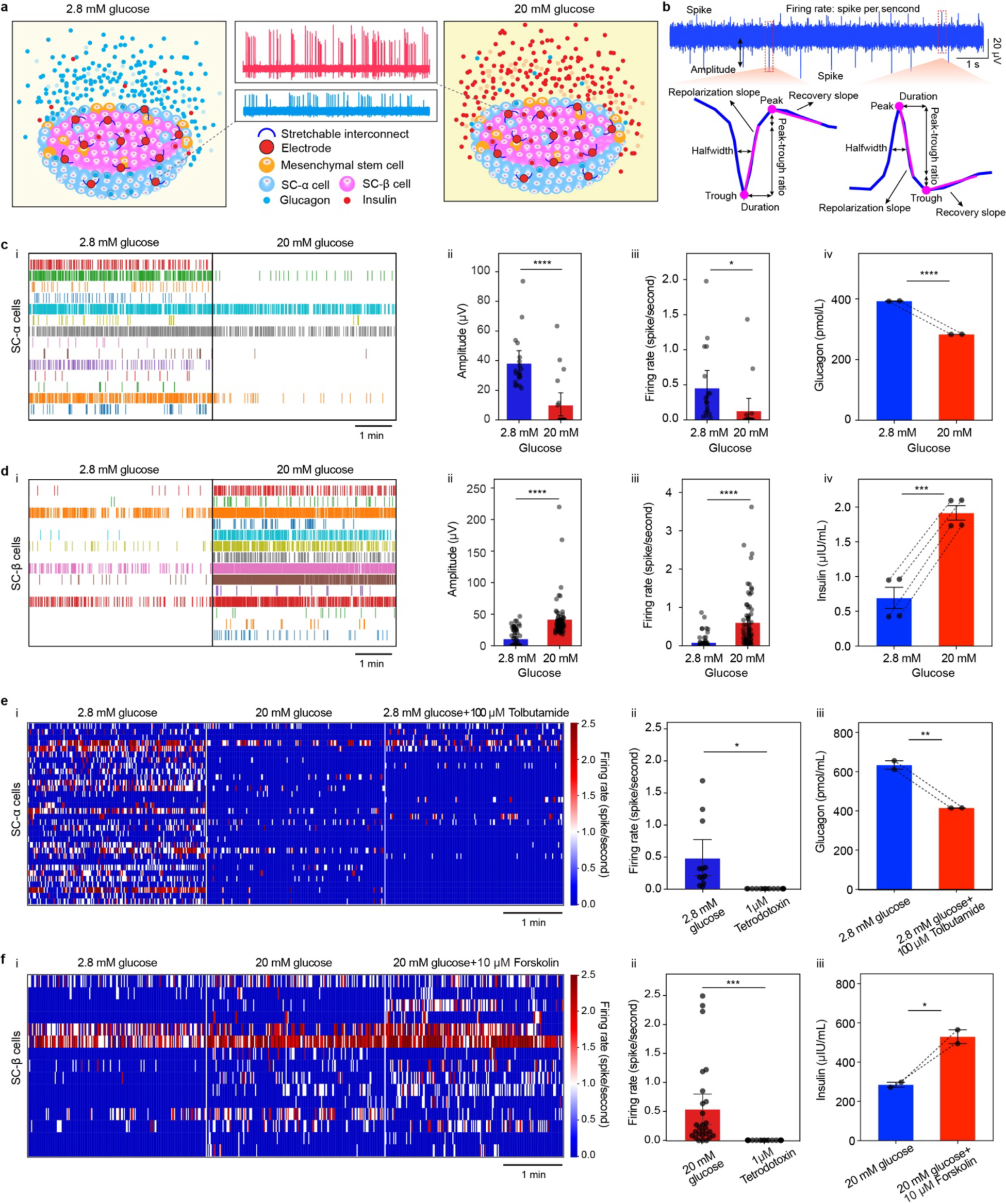
Simultaneous tracing of islet-wide cell type-specific and stimulus-coupled electrical dynamics. **a,** Schematics showing SC-α and SC-β cells present electrical activities, voltage-gated glucagon and insulin secretion in response to low (2.8 mM) and high (20 mM) glucose stimulation, respectively, upon functional maturation. **b,** Cyborg islet cell type-specific electrical features analyzed. Spike firing rate, amplitude, duration, peak-trough ratio, width at half-maximum, repolarization slope, and recovery slope were compared between 2.8 mM and 20 mM glucose stimulation. **c**, SC-α cell spike trains (i), average amplitude (ii), and average firing rates (iii) recorded at 2.8 mM and 20 mM glucose (bar plots show mean ± SEM of N = 19 cells pooled from n = 5 cyborg islets). Glucagon secretion into the medium measured by protein ELISA at 2.8 mM and 20 mM glucose (iv) (data are mean ± SEM from n = 2 replicate measurements). **d**, SC-β cell spike trains (i), average amplitude (ii), and average firing rate (iii) recorded at 2.8 mM and 20 mM glucose (bar plots show mean ± SEM of N = 71 cells pooled from n = 5 cyborg islets). Insulin secretion into the medium measured by protein ELISA at 2.8 mM and 20 mM glucose (iv) (data are mean ± SEM from n = 2 replicate measurements). **e**, Heatmap showing firing rate of SC-α cells recorded at 2.8 mM glucose, 20 mM glucose, and 2.8 mM glucose + 100 μM tolbutamide (i) (N = 32 cells pooled from n = 3 cyborg islets). Firing rate per minute of SC-α cells recorded at 2.8 mM glucose and 1 μM tetrodotoxin (ii) (bar plots show mean ± SEM of n = 12 pooled from n = 2 cyborg islets). Glucagon secretion into the medium measured by protein ELISA at 2.8 mM glucose and 2.8 mM glucose + 100 μM tolbutamide (iii) (data are mean ± SEM from n = 2 replicate measurements). **f**, Heatmap showing firing rate of SC-β cells recorded at 2.8 mM glucose, 20 mM glucose, and 20 mM glucose + 10 μM forskolin (i) (N = 15 cells pooled from n = 3 cyborg islets). Firing rate per minute of SC-β cells recorded at 20 mM glucose and 1 μM tetrodotoxin (ii) (bar plots show mean ± SEM of n = 27 pooled from n = 2 cyborg islets). Insulin secretion into the medium measured by protein ELISA at 20 mM glucose and 20 mM glucose + 10 μM forskolin (iv) (data are mean ± SEM from n = 2 replicate measurements). n.s., not significant; * p < 0.05; ** p < 0.01; *** p < 0.001; **** p < 0.0001 from two-tailed, unpaired t tests.

We observed two distinct clusters of cells exhibiting differential electrical activity in response to low/high glucose. One cluster showed greater activity under 2.8 mM glucose (Fig. 2c), while the other was more active under 20 mM glucose (Fig. 2d). Specifically, spike firing was elevated under 2.8 mM relative to 20 mM glucose for cells in the first cluster (Fig. 2ci), with significantly higher average amplitude and firing rate (Fig. 2cii-iii). Glucagon secretion into the medium was also higher under 2.8 mM glucose (Fig. 2civ). The second cluster, by contrast, displayed greater spike firing under 20 mM compared to 2.8 mM glucose (Fig. 2di), with significantly increased average amplitude and firing rate (Fig. 2dii-iii). Moreover, insulin secretion into the medium was significantly higher under 20 mM glucose (Fig. 2div). Based on the known glucose-responsive behaviors of α and β cells^8,37^, we classified the first cluster as α-like (SC-α) cells and the second as β-like (SC-β) cells.

To validate the classification and cell type-specific electrical activities of SC-α and SC-β cells, we next measured electrical responses to additional secretion modulators, including tolbutamide (K_ATP_ channel inhibitor)^36,38^, forskolin (cyclic adenosine monophosphate / protein kinase A [cAMP/PKA] activator)^15^, and tetrodotoxin (TTX, Na^+^ channel blocker)^39^ (Fig. 2e, f). We first assigned SC-α and SC-β cells based on electrical activities in 2.8 mM and 20 mM glucose, and then examined cell type-specific responses to specific drugs. For SC-α cells, we observed decreased firing rates under tolbutamide compared to 2.8 mM glucose alone (Fig. 2ei, Extended Data Fig. 3c). As expected, glucagon secretion was lower under tolbutamide relative to 2.8 mM glucose alone (Fig. 2iii), consistent with tolbutamide’s inhibitory effect on glucagon release in α cells^36^. In contrast, for SC-β cells, we saw an increase in firing rate under forskolin relative to 20 mM glucose alone (Fig. 2fi, Extended Data Fig. 3d). Insulin secretion was higher under forskolin compared to 20 mM glucose alone (Fig. 2fiii), consistent with forskolin enhancing electrical activity and insulin secretion in β cells^15^. Finally, we exposed cyborg SC-islets to TTX, which effectively inhibited electrical activity in both stimulated SC-α and β cells (Fig. 2eii,fii). In summary, using the cyborg SC-islet platform, we measured simultaneous α-like and β-like specific electrical activities and validated their distinct responses to glucose and secretion modulators.

### Tracking maturing SC-α and β electrical activities

Cyborg SC-islets offer a unique platform to trace islet-wide SC-α and β electrical activities and their stimulus-coupled dynamics as they evolve during functional maturation. Using embedded electrodes, we monitored glucose-stimulated electrical activities during extended culture from weeks 2 to 7 after device integration (Fig. 3a). The results show that electrical signals significantly increase after 2 weeks of extended culture, with progressively greater numbers of recorded cells acquiring electrophysiological function (Fig. 3a). We then used our spike sorting approach to trace the electrical maturation trends of individual SC-α and SC-β cells. Clustering cell-level firing rates throughout the extended culture time course identified SC-α and β cells, based on greater firing under 2.8 mM or 20 mM glucose, respectively, and other cells with unchanging / no electrical activity under 2.8 mM or 20 mM glucose (Fig. 3b, Extended Data Fig. 4aiii).

**Fig. 3.**
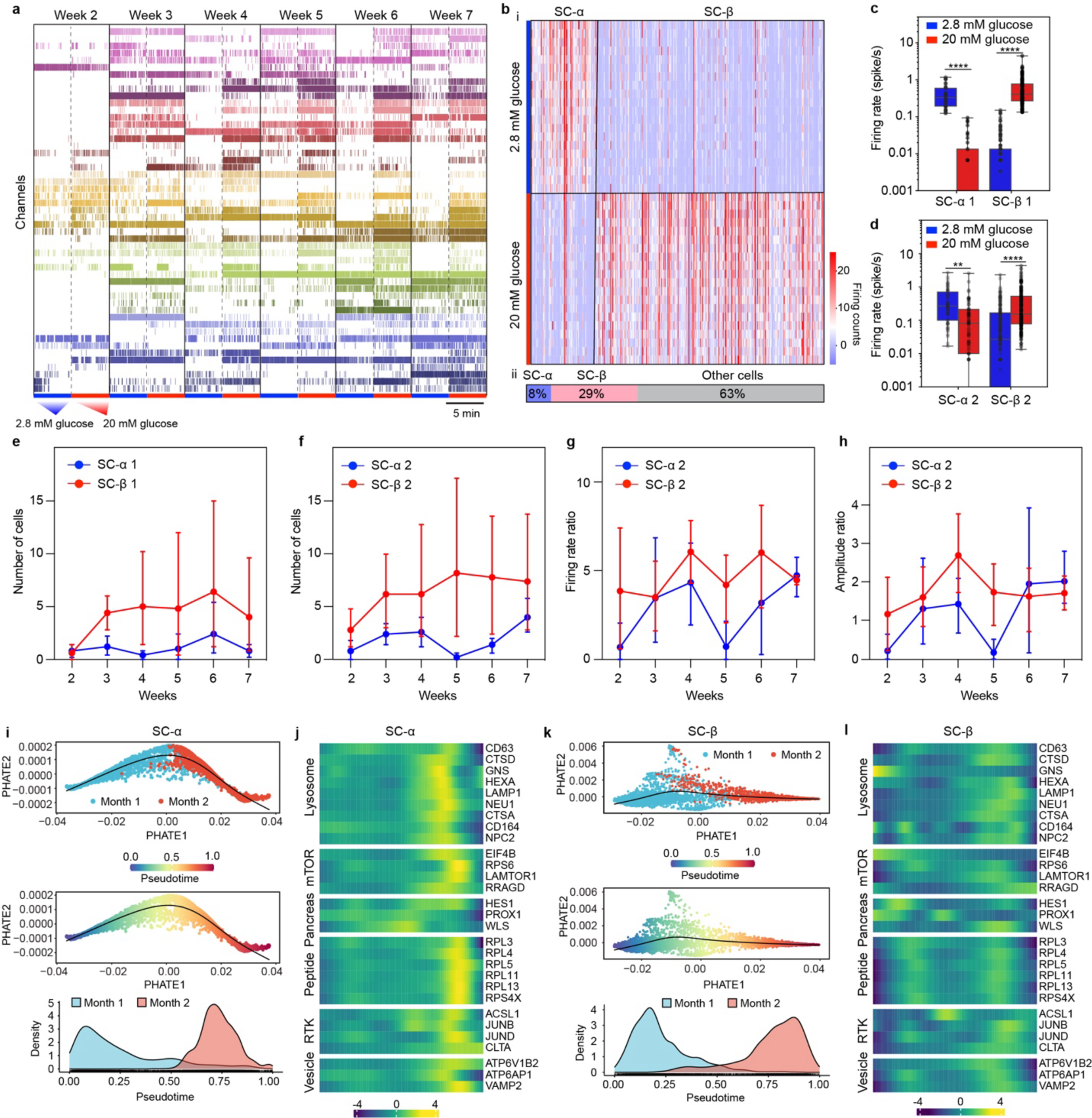
Long-term stable tracking of SC-α and SC-β electrical activities during early functional maturation. **a**, Representative electrical spike trains recorded over 5 minutes at 2.8 mM and 20 mM glucose from weeks 2 to 7 after cyborg islet device integration. Cyborg islets were cultured in serum-free medium without exogenous signaling factors. Data was collected from 6 cyborg islets. **b**, Clustering of firing rates during extended culture of cyborg islets identifies cells with increased firing under 2.8 mM or 20 mM glucose as SC-α or SC-β, respectively (i). The percentage of SC-α, SC-β, and other cells (ii, See Methods) **c,** Spike firing rates for SC-α 1 and SC-β 1 cells at 2.8 mM and 20 mM glucose during extended culture after device integration (bar plots show mean ± SEM of N = 33 SC-α 1 and 126 SC-β 1 cells pooled from n = 5 cyborg islets). **d,** Spike firing rates for SC-α 2 and SC-β 2 cells at 2.8 mM and 20 mM glucose during extended culture after device integration (bar plots show mean ± SEM of N = 57 SC-α 2 and 193 SC-β 2 cells pooled from n = 5 cyborg islets). **e,** The number of electrically recorded SC-α 1 and SC-β 1 cells from week 2 to week 7 after device integration. Data are mean ± SEM of N = 33 SC-α 1 and 126 SC-β 1 cells pooled from n = 5 cyborg islets. **f,** The number of electrically recorded SC-α 2 and SC-β 2 cells from week 2 to week 7 after device integration. Data are mean ± SEM of N = 57 SC-α 2 and 193 SC-β 2 cells pooled from n = 5 cyborg islets. **g,** Spike firing rate ratios for SC-α 2 (2.8 mM / 20 mM glucose) and SC-β 2 (20 mM / 2.8 mM glucose) cells from week 2 to week 7 after device integration. Data are mean ± SEM of N = 48 SC-α 2 and 126 SC-β 2cells pooled from n = 5 cyborg islets. **h,** Spike amplitude ratios for SC-α 2 (2.8 mM / 20 mM glucose) and SC-β 2 (20 mM / 2.8 mM glucose) cells from week 2 to week 7 after device integration. Data are mean ± SEM of N = 48 SC-α 2 and 126 SC-β 2 cells pooled from n = 5 cyborg islets. **i**, Early maturation trajectory of SC-α cells using 2D PHATE visualization, by month (top) or inferred pseudotime ordering (middle), and pseudotime distributions of month 1 and month 2 islet cells (bottom). **j**, SC-α gene expression programs along with their enriched pathways. Heatmap shows the row z-scored expression of selected dynamic genes along SC-α pseudotime. **k**, Early maturation trajectory of SC-β cells using 2D PHATE visualization, by month (top) or inferred pseudotime ordering (middle), and pseudotime distributions of month 1 and month 2 islet cells (bottom). **l**, SC-β gene expression programs along with their enriched pathways. Heatmap shows the row z-scored expression of selected dynamic genes along SC-β pseudotime.

By comparing average spike firing rates and amplitudes under 2.8 mM and 20 mM glucose, we identified two major states for SC-α and β cells, which can be further classified into SC-α1/α2 and SC-β1/β2 cell states (Extended Data Fig. 4ai,ii, see Methods). Specifically, SC-α1/β1 cells show low spike firing rate (typically <0.1 spikes/second) and amplitude (typically <10 μV) under non-stimulatory 20 mM/2.8 mM glucose, respectively (Fig. 3c, Extended Data Fig. 4b-e). These low basal firing rates increase significantly when SC-α1/β1 cells are respectively exposed to stimulatory 2.8 mM/20 mM glucose. Conversely, SC-α2/β2 cells exhibit higher basal firing rate and amplitude at 20 mM/2.8 mM glucose, respectively, with comparable elevation of spike firing rates under stimulatory 2.8 mM/20 mM glucose (Fig. 3d, Extended Data Fig. 4b-e). Moreover, cells in SC-α1/β1 and SC-α2/β2 states increased from week 2 to week 7 in all recorded SC-islets (Fig. 3e,f). As a metric of stimulation capacity, we calculated the ratio of spike firing rates/amplitudes between stimulatory and basal glucose conditions (2.8 mM/20 mM glucose for SC-α, and 20 mM/2.8 mM glucose for SC-β). We find that firing rate and amplitude ratios for both α2 and β2 cells gradually increased from week 2 to week 7 (Fig. 3g,h). To further delineate maturing SC-α1/α2 and SC-β1/β2 electrical activities, we analyzed their duration, peak-trough ratio, width at half-maximum, repolarization slope, and recovery slope. We found no significant time-dependent changes in these waveform features over the extended culture time course under low/high glucose, indicating that extracellular spikes remain largely unchanged during *in vitro* maturation (Extended Data Fig. 4f-o). Our longitudinal electrical recording studies reveal the following findings: First, SC-α and β cells can be further subclassified by their basal activities. SC-α2 and β2 cells fire more under non-stimulatory glucose, as with immature (fetal/neonatal) cells, which display a low glucose threshold for action potential firing^40^ and secretion^16^. Second, the turnover of SC-α and β cells in these states is highly dynamic over a 2-month extended culture period, with increasing numbers of cells entering both less and more mature states. Interestingly, despite these changes, spike waveform features of both SC-α and β cells remain relatively stable during long-term culture. Third, cells in less mature states (SC-α2 and β2) show gradually increasing glucose stimulation capacity during the culture period, indicating ongoing specialization of glucose responsiveness. In sum, our findings reveal population-level changes in maturing islet cell states over time, as well as cell-level changes in stimulus-coupled electrical responses during maturation.

To investigate gene regulatory events underlying electrical maturation during extended culture, we performed scRNA-seq on cyborg islets at months 1 and 2 after device integration (Fig. 3i-l, Extended Data Fig. 5). Specifically, we conducted pseudotime analysis to reconstruct continuous early maturation trajectories for SC-α, β, and EC cells. We then used a two-step analysis to elucidate transcriptional changes along their maturation trajectories. First, we projected expression profiles using PHATE^41^ followed by pseudotime inference using Slingshot^42^ (Fig. 3i,k). We then analyzed enriched functions among genes upregulated along pseudotime, illuminating the transcriptional maturation process of the endocrine subtypes (Extended Data Fig. 5a,b). This analysis revealed a multifaceted maturational landscape. Ribosomal components (e.g., *RPL3*, *RPL4*, *RPL5*) and targets of receptor tyrosine kinase signaling (e.g., *JUNB*, *JUND*, *CLTA*) were enriched across all cell types. Mediators of lysosomal activity (e.g., *CD63*, *CTSD*, *LAMP1*) and ATP-driven vesicle budding (e.g., *VAMP2*, *ATP6AP1*) were enriched within SC-α and β cells, consistent with their coupling of energy metabolism to hormone secretion (Fig. 3j, l). Conversely, mTOR pathway-associated genes associated with nutrient sensing and energy balance (e.g., *EIF4B*, *LAMTOR1*) were specifically upregulated in SC-α cells (Fig. 3j). These results link energy and hormone metabolism gene pathways to islet cell type-specific maturation trajectories, implicating them in the coupling and optimization of electrical/secretory activities to nutrient sensing.

### Tracing maturation triggered by daily feeding cycles

Entrainment to 24-hour feeding-fasting cycles enhances *in vitro* maturity of SC-islets^8^. However, the specific role of circadian rhythms on electrical maturation of SC-α and β cells has not been investigated. After 2-month extended culture, we thus subjected cyborg islets to 24-hour glucose and forskolin shock and recovery cycles over a four-day timeframe, followed by electrical recording in a constant environment, to dissect the impact on SC-islet electrical maturation (Fig. 4a). Immunostaining of cyborg islets shows that effective coupling of the nanoelectronics with SC-α and β cells is retained at 2 months after device integration with/without circadian entrainment (Extended Data Fig. 6a,b). Following circadian entrainment, we conducted electrical measurements at 2.8 mM and 20 mM glucose every 4 hours for 72 hours, thereby capturing hourly electrical trends (Fig. 4b,c). In cyborg islets without entrainment, a gradual islet-level loss of electrical activity was evident over the three days of recording, whereas circadian-entrained samples exhibited increasing electrical activity (Extended Data Fig. 6c,d). These results evidence how diurnal feeding-fasting within a relatively short period of four days enhances overall *in vitro* SC-islet function.

**Fig. 4.**
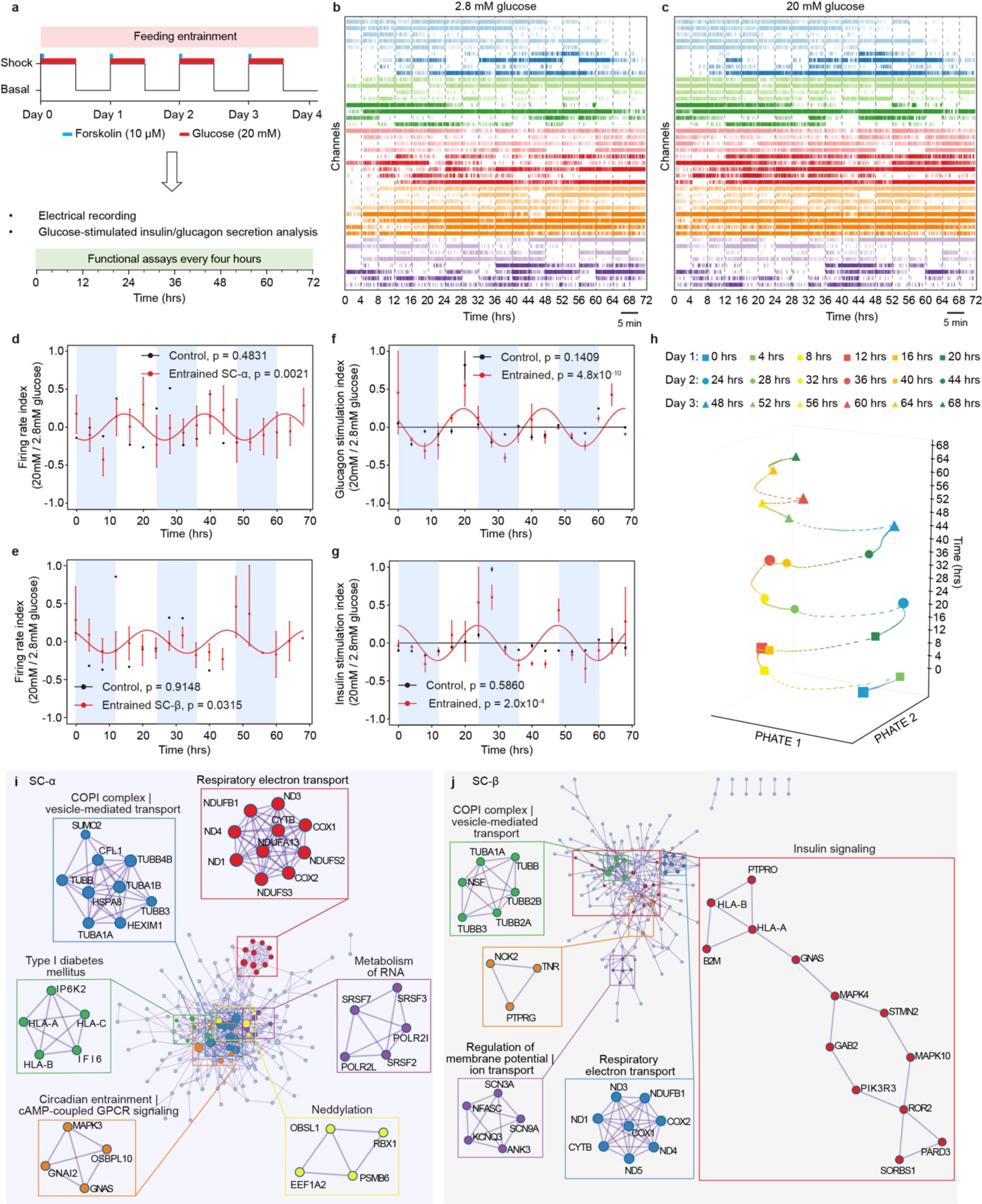
Tracing of SC-α and SC-β electrical maturation triggered by circadian entrainment. **a**, Timeline for metabolic shock/recovery cycles conducted 4 times over 4 days, followed by functional assays, including electrical and insulin/glucagon secretion measurements at 2.8 mM and 20 mM glucose, conducted every 4 hours for 72 hours. **b-c**, Spike trains recorded over 5 minutes at 2.8 mM (**b**) and 20 mM (**c**) glucose every 4 hours for 3 days following circadian entrainment. **d-e**, Spike firing rate ratios for SC-α (**d**) and SC-β (**e**) cells recorded over 72 hours after cyborg islet diurnal feeding (entrained) or mock (control) entrainment. Data are detrended (baseline-subtracted) mean ± SEM of N = 210-420 SC-α and SC-β cells pooled from 5 independent entrained organoids and of N = 60-52 SC-α and SC-β cells from a control islet. **f-g**, Glucose stimulation indexes for glucagon (**f**) and insulin (**g**) secretion into the medium measured by protein ELISA over 72 hours after cyborg islet diurnal feeding (entrained) or mock (control) entrainment. Data are detrended (baseline-subtracted) mean ± SEM of 5 independent entrained cyborg islets and a control islet, with n = 2 replicate measurements each. **h**, 3D PHATE visualization of the averaged spike waveforms of N = 341 SC-β cells over 72 hours at 20 mM glucose from an entrained cyborg islet. **i-j**, Biological functions enriched among protein-protein interaction complexes identified by the MCODE^60^ algorithm based on genes upregulated in SC-α (**i**) and SC-β (**j**) cells in circadian-entrained versus control cyborg islets. For SC-α cells, the top 6 interaction complexes identified (including COPI complex | vesicle-mediated transport, respiratory electron transport, type 1 diabetes mellitus, metabolism of RNA, and circadian entrainment | cAMP-coupled GPCR signaling) are highlighted (**i**); For SC-β cells, the top 5 interaction complexes identified (including insulin signaling, COPI complex | vesicle-mediated transport, respiratory electron transport, and regulation of membrane potential | ion transport) are highlighted (**j**).

Following entrainment, cell-level electrical traces revealed coordinate circadian rhythms in glucose-coupled spike firing rates within individual SC-α and β cells (Fig. 4d,e, Extended Data Fig. 6f,h). By contrast, SC-α and β cells did not display rhythmic firing without entrainment, and lost electrical activity after 2 days of recording (Extended Data Fig. 6e,g). Importantly, glucagon and insulin secreted into the medium of entrained samples showed clear circadian oscillation, while control samples did not exhibit rhythmic hormone secretion (Fig. 4f,g). A phase shift was observed between glucagon and insulin glucose responsiveness rhythms, as with previous *in vitro* studies^43^, which was reflected at the level of SC-α and β cell electrical activities (Fig. 4d-g, Extended Data Fig. 6f,h). To further investigate circadian electrical activity, we used PHATE^41^ to project the time evolution of averaged spike waveforms of SC-β cells under 20 mM glucose from an entrained cyborg islet over the 72h of recording (Fig. 4h), revealing time-of-day variation in waveform profiles. These results indicate that diurnal feeding-fasting entrains population and cell-level electrical activities of both α-like and β-like cells within SC-islets to elicit sustained circadian secretory responses to glucose.

To investigate genetic mechanisms underlying enhanced electrical function upon circadian entrainment, we compared scRNA-seq from entrained cyborg islets relative to parallel unentrained ones (Fig. 4i,j, Extended Data Fig. 6k-o). Genes for respiratory electron transport machinery were consistently induced across SC-α, β, and EC cells, with NADH dehydrogenase (e.g., *ND1-5*) and cytochrome b/c (e.g., *CYTB*, *COX1-2*) components identified as the top protein-protein interaction complex in each endocrine subtype. Vesicular transport machinery (e.g., tubulins) was mainly upregulated within SC-α and β cells, with COPI coat proteins mediating vesicle budding identified as a top complex in these cells, consistent with their secretory functions. Insulin signaling (e.g., MAP kinases) and HLA (human leukocyte antigen) genes associated with type I diabetes were also induced in entrained SC-α and β cells. Ion channels play a key role in cellular electrophysiology, with many genes being upregulated upon entrainment that directly regulate hormone release. Notably, entrainment induced subunits of voltage-gated K^+^ (e.g., *KCNQ3*), Na^+^ (e.g., *SCN3A/9A*), and Ca^2+^ (e.g., *CACNA1E, CACNG4*) channels, as well as ATP-sensitive K^+^ channels (e.g. *ABCC8, ABCC9*), that regulate membrane potential in SC-β cells (Fig. 4j, Extended Data Fig. 6l-o); interacting cAMP-coupled GPCR signaling components (e.g., *GNAS, GNAI2*) linked to circadian entrainment in SC-α cells (Fig. 4i, Extended Data Fig. 6l-n); and Ca^2+^ transmembrane transporters in SC-EC cells (Extended Data Fig. 6k-n). Finally, we observed upregulation of cell-cell communication genes across all cell types, including various cadherins and cell adhesion molecules that form cell-cell junctions, consistent with coordinate SC-α and β electrical activities. Together, our results reveal the following findings: First, daily feeding modulation reprograms glucose responsiveness at multiple steps, from glucose oxidation to Ca^2+^-dependent hormone vesicle release. Interestingly, entrainment induces key islet developmental controllers (e.g., *ISL1, PAX6, ARX, WNT4*) that may act as upstream regulators of these diverse pathways. Second, sustained electrical activity upon circadian entrainment involves upregulation of ion transporters that directly regulate membrane potential. Third, synchronization of entrained α/β cells is linked to greater cell-cell communication, potentially contributing to greater coordination of their stimulated activities. In sum, our findings reveal that feeding-fasting synchronizes islet-wide α-β activities to generate coordinated stimulus-coupled hormone secretion rhythms via cell-cell communication and exocytic network induction.

## Discussion

How α and β cell activities within human islets evolve and become coordinated during the critical period of functional maturation, when they reach full secretory capacity, has remained largely unexplored, either in native or *in vitro* settings. Here, we introduce a novel approach to investigate the physiological maturation of human SC-islets using embedded electrodes, allowing capture of the membrane potential changes that underlie hormone release. This approach offers a unique opportunity for 1) real-time electrophysiological recording of simultaneous α-like and β-like cell activities across the entire islet and 2) continuous tracking of islet-wide single-cell activities throughout the maturation process. These capabilities open new avenues for discovery-based studies of changes in islet physiology underlying maturation. Moreover, they offer quantitative readouts for genetic and drug screenings aimed at identifying maturation-modulating factors, paving the way for hypothesis-driven studies. Employing our cyborg islet platform, we characterized electrical activities of α-like and β-like cells within SC-islets as their function specializes during extended *in vitro* culture and upon circadian entrainment. Our data reveal several new findings. First, SC-α and β cells in more mature states, marked by low basal firing rates, can be found from the earliest maturation stages. Second, a higher islet-level glucose threshold for hormone release during maturation involves transitions in cell-level electrical activities. Previous work linked the increased glucose threshold for insulin secretion during β cell maturation to greater Ca^2+^ sensitivity of insulin vesicles^44^. Our studies indicate that membrane depolarization, which occurs upstream and triggers extracellular Ca^2+^ influx, also plays a role. Third, circadian rhythms enhance coordinated stimulus-coupled hormone responses by synchronizing islet-wide α and β electrical activities and waveform profiles. These findings expand our understanding of islet physiology, complementing observations from calcium imaging and hormone secretion studies about β-cell maturation^45^ and heterogeneity^46^. Moreover, they offer an explanation for how circadian rhythms lead to improved islet function^8^.

Our studies also delineate gene regulatory events underlying stepwise α-like and β-like cell maturation in SC-islets. During early maturation, prompted by extended culture, we see gradual induction of ribosomal genes across endocrine cells, consistent with increasing secretory burden, as observed previously^12,14^. We also see dynamic modulation of mTOR and lysosomal pathway-associated genes linked to nutrient sensing and energy balance, consistent with increased coupling of hormone secretion to energy metabolism^13^. During further *in vitro* maturation, prompted by daily feeding-fasting, we reveal dynamic regulation of electrophysiology-modulating genes governing hormone secretion in SC-α and β cells. These include key ion channel components, such as *ABCC8*, encoding the SUR1 subunit of the ATP-sensitive K^+^ channel^47^; *SCN3A* and *SCN9A*, encoding the primary physiologically relevant α and β cell Na^+^ channels, respectively^48^; and *CACNA1G* and *CACNA1E*, encoding subunits of voltage-gated Ca^2+^ channels that modulate calcium currents vital for β-cell electrophysiology^46,49^. We also see induction of oxidative respiration and hormone metabolism genes, including electron transport and exocytic machinery, evidencing refinement of glucose responsiveness at multiple levels (metabolic, electrical, Ca^2+^ handling, and secretory capacity). Interestingly, cell-cell junction and HLA genes are also induced during late maturation, suggesting that circadian synchronicity of endocrine cell activites involves enhanced cell-cell communication and impacts antigen presentation. These insights hold potential not only for research applications, such as dissecting the interplay between maturity and immunogenicity, but also for the design of fully functional cell-based therapeutics for autoimmune diabetes.

Further delineating mechanisms for the evolution, propagation, and synchronization of secretion dynamics in SC-islets will require paired mapping of cellular activities and gene expression, as islets rely on spatiotemporally orchestrated interactions among heterogenous cells to sustain specialized functions. Future work will thus incorporate our recently developed *in situ* electro-sequencing platform^27^ to spatially resolve cellular identity and functionality during islet maturation. Although we demonstrate the integration of sensors within the cyborg islet platform, we anticipate that future advancements in this direction will enable multimodal characterizations, such as metabolite and oxygen sensing^50^. Leveraging the recently demonstrated ability of electrical stimulation to promote maturation of stem cell-derived organoids^51,52^ will in turn enable feedback control, such as tunable electrical or chemical stimulation^50^. Thus, we expect that future progress in multifunctional stretchable mesh nanoelectronics will allow for closed-loop sensing and actuation, offering a powerful approach to further improve organoid function.

## Methods

### Device fabrication, assembly, and characterization

Stretchable mesh nanoelectronics were fabricated following previously established methods^24–27^. First, a 500-μm-thick glass wafer was cleaned with acetone, isopropyl alcohol, and deionized (DI) water. Next, a 100-nm-thick nickel (Ni) sacrificial layer was deposited using a thermal evaporator (Sharon). The SU-8 precursor (SU-8 2000.5, MicroChem) was spin-coated to achieve a thickness of either 800 or 400 nm, followed by pre-baking at 65°C and 95°C for 2 min each. The SU-8 was then exposed to 365 nm UV for 200 mJ/cm^2^, post-baked at 65 °C and 95 °C for 2 min each, developed using SU-8 developer (MicroChem) for 60 s, and baked at 180°C for 40 min to define mesh-like SU-8 patterns for bottom encapsulation. Then, the LOR3A photoresist (MicroChem) was spin-coated at 4000 rpm, pre-baked at 180°C for 5 min, and the S1805 photoresist (MicroChem) was spin-coated at 4000 rpm, pre-baked at 115°C for 1 min, exposed to 405 nm UV for 40 mJ/cm^2^, and developed using CD-26 developer (Micropost) for 70 s to define interconnect patterns. Chromium/gold/chromium (Cr/Au/Cr) with a thickness of 5/40/5 nm was deposited using an electron-beam evaporator (Denton), followed by a standard lift-off procedure in remover PG (MicroChem) to define the interconnects. Electrode patterns were defined in the LOR3A/S1805 bilayer photoresists as described above. Chromium/platinum (Cr/Pt) with a thickness of 5/50 nm was deposited using an electron-beam evaporator (Denton), followed by a standard lift-off procedure in remover PG (MicroChem) to define the electrodes. Finally, the top SU-8 encapsulation layer was defined as described above.

Multi-channel flexible flat cables (FFCs, Molex) were soldered onto the input/output (I/O) pads using a flip-chip bonder (Finetech Fineplacer). Subsequently, a custom-made cell culture chamber was affixed to the substrate wafer with a biocompatible adhesive (Kwik-Sil, WPI). To achieve Pt black electroplating on the Pt electrode array, a 0.08 wt% chloroplatinic acid (H_2_PtCl_6_) precursor solution in DI water was drop-casted onto the device. A direct current (DC) electrical current density of 1 mA/cm^2^ was then applied for 3 min using the electrodes as anodes and an external Pt wire as the cathode. The device was thoroughly rinsed with DI water and dried with N_2_ before undergoing treatment with oxygen plasma (Anatech 106 oxygen plasma barrel asher). Finally, a 1 mL Ni etchant (type TFG, Transene) was introduced into the chamber for 2 to 4 h to completely release the stretchable mesh nanoelectronics from the glass substrate.

To determine the electrochemical impedance spectrum of electrodes in each device, we employed a three-electrode setup. The counter electrode comprised a platinum wire (300 μm in diameter, 1.5 cm in length immersed), while a standard silver/silver chloride electrode served as the reference electrode. For the measurements, we utilized the SP-150 potentiostat (BioLogic) and its commercial software EC-lab. To establish statistical significance, we performed at least three frequency sweeps for each measurement, ranging from 1 MHz to 1 Hz. A sinusoidal voltage of 100 mV peak-to-peak was applied, and the response to ten consecutive sinusoids (spaced out by 10% of the period duration) was accumulated and averaged for each data point. Additionally, crosstalk between electrodes was evaluated at 1 kHz using a Blackrock CerePlex Direct voltage amplifier. To perform cell culture, the devices were rinsed in DI water three times, followed by sterilization through immersion in 70% ethanol for 15 min. Then, the devices were washed with DPBS and incubated with Poly-D-lysine hydrobromide (0.01% w/v) and Matrigel solution (100 μg/mL). Finally, 60 μL liquid Matrigel (10 mg/mL) (hESC-Qualified Matrix, Corning) was added to the cell culture chamber to form a Matrigel hydrogel substrate before cell culture.

### Cell culture and differentiation

Human embryonic pluripotent stem cells (HUES8 NIH hESC registry #09-0021) were seeded at a density of 0.6 million/ml in mTeSR1 medium (StemCell Technologies) supplemented with 10 μM ROCK inhibitor Y27632 (DNSK International) for directed differentiation toward islet organoids as described previously^7,8,30^. After seeding, a half-feed with mTeSR1 was performed after 24 h, followed by a full feed with mTeSR1 after 48 h. At 72 h post-seeding, the following stepwise differentiation protocol was initiated (culture media and differentiation factors are as described previously^7,8,30^:

Stage 1 (3 days in S1 medium):

- Day 1: 100 ng/mL ActivinA + 14 μg/mL CHIR99201
- Day 2: 100 ng/mL ActivinA
- Day 3: no media change.

Stage 2 (3 days in S2 medium) with 50 ng/mL KGF, feeding every other day.

Stage 3 (2 days in S3 medium):

- Day 1: 50 ng/ml KGF + 0.25 μM Sant1 + 2 μM Retinoic acid (RA) + 500 nM PDBU + 10 μM ROCK inhibitor + 200 nM LDN193189
- Day 2: Same factors as Day 1, except LDN193189 is omitted.

Stage 4 (5 days in S3 medium) with 50 ng/ml KGF + 0.25 μM Sant1 + 0.1 μM Retinoic acid (RA) + 10 μM ROCK inhibitor + 5 ng/mL Activin A, feeding every other day.

Stage 5 (7 days in BE5 medium):

- Day 1-4: 0.25 nM Sant1 + 20 ng/mL Βetacellulin + 1 μM XXi + 10 μM Alk5i II + 1 μM T3 + 0.1 μM RA. Media changed every other day.
- Day 5-7: Media containing no Sant 1 and 0.025 μM RA. Media changed every other day.

Stage 6: Extended culture in S3 medium, with feeding every other day.

Human mesenchymal stem cells (hMSCs; PT-2501) obtained from Lonza (Walkersville, MD, USA) were maintained in 6-well plates using MSCGM BulletKit medium (cat # PT-3238 & PT-4105, Lonza).

All methods involving human cells were approved by the Harvard University and University of Pennsylvania IRB and ESCRO committees.

### Cyborg human SC-islet assembly

Human SC-islet organoids were dissociated into a single cell suspension as described previously^8,30^. Briefly, upon completion of differentiation, organoids were collected in suspension medium, washed with an equal volume of DPBS, and then incubated in 6.5 mL DPBS and 8 mL Accutase (StemCell Technologies) for 7 min at room temperature. After washing, the organoids were dissociated into single cells in PBS + 10 μM ROCK inhibitor by mechanical pipetting up and down 40-50 times with a P1000 set to 1 mL. For MSC dissociation, trypsin-EDTA 0.05% was used, and trypan blue was used to count the cells. To integrate cells with stretchable mesh nanoelectronics, SC-islet single cells (1×10^6^ cells per 16-channel device and 4×10^6^ cells per 64-channel device) and hMSCs (0.5×10^5^ cells per device) were suspended in a mixture (3:1) of S3 medium and MSCGM medium. The cell mixture was then transferred into the device-containing cell culture chamber as described above and cultured at 37°C with 5% CO_2_.

### *In vitro* circadian entrainment

Cyborg human SC-islets were subjected to circadian entrainment following a previously described method^8^. Specifically, 24-hour metabolic shock / recovery cycles were applied as follows: cells were cultured in S3 medium containing 20 mM Glucose (Sigma; G7528) + 10 μM Forskolin (Stemgent; 04-0025) for 1 h, then washed and cultured in S3 medium containing 20 mM Glucose for the remaining 11 h. Afterward, they were washed and cultured in basal S3 medium for a 12-hour recovery period. The shock / recovery treatments were repeated four times, with a total duration of four days. Stringent washes were performed between media switches to ensure complete removal of any remaining added factors. The same washing and incubation times were used for control samples mock-treated with basal S3 medium only.

### Electrical recordings

Electrical activity was recorded using a RHD 64-channel headstage (Intan technologies) connected to the Intan 1024 ch recording controller (Intan technologies). A Pt electrode was employed for grounding the culture medium, while another Pt electrode served as the reference electrode. The samples were placed on a battery-powered warming plate to maintain a thermostatic 37 °C during electrical measurements. The entire measurement setup was enclosed in a Faraday cage. The electrical recording was performed at a sampling rate of 20,000 Hz. For long-term electrical recordings, the samples were recorded for 5 mins weekly at both 2.8 mM and 20 mM glucose concentrations. For samples subjected to circadian entrainment, upon completion of metabolic shock/recovery cycles, electrical recordings were performed every 4 h for 72 h at 2.8 mM and 20 mM glucose concentrations.

### Glucose-stimulated insulin/glucagon secretion assays

Cyborg human SC-islets were washed twice with Krebs buffer containing 2.8 mM glucose, followed by a one-hour incubation in 2.8 mM glucose Krebs buffer to remove residual insulin. Next, samples were washed with 2.8 mM glucose Krebs buffer, and sequentially exposed to Krebs buffer containing 2.8 mM glucose and 20 mM glucose, with a one-hour incubation time for each concentration. An additional wash was carried out between the 2.8 mM and 20 mM glucose incubations to remove residual glucose. All incubations were conducted at 37 °C, and supernatant samples were collected at the end of each incubation. The levels of human insulin and glucagon in the collected supernatants were determined using a Human Ultrasensitive Insulin ELISA KIT (ALPCO Diagnostics; 80-INSHUU-E10) and a Glucagon ELISA KIT (Mercodia; 10-1271-01), respectively, according to the manufacturer’s instructions. Briefly, all collected supernatants were thawed and mixed well before use, and duplicate 25 μL samplings were assayed to ensure reliability of measurements. Samplings were mixed with detection antibodies in Kit-provided 96-well plates, which were then sealed and incubated at 750 rpm on a plate shaker at the indicated temperatures and incubation times. Unbound antibodies were removed by washing plates 6 times avoiding the formation of bubbles and ensuring removal of all buffers from the last wash. Bound conjugates were detected by incubation at 750 rpm with 3,3’,5,5’-tetramethylbenzidine (TMB), which was allowed to proceed for the indicated incubation time (or for a shorter time if the color of the more concentrated Kit insulin/glucagon concentration standard was strong enough). Reactions were then stopped by adding Kit-provided acidic stop solution to the plate, which was placed on a plate shaker for about 10 s to homogenize reactions. Colorimetric endpoints were read in a CLARIOstar microplate reader (BMG Labtech) using 450 nm excitation, and insulin/glucagon concentration was quantified for each sample based on the Kit-provided concentration standards.

### Immunostaining and imaging

Cyborg and control SC-islets were subjected to immunostaining and clearing procedures as previously described^24,27^. The primary antibodies used in the staining process included Rat anti-INS (Cat# GN-ID4, RRID: AB_2255626, DSHB, 1:100); Mouse anti-GCG (SC-514592, Santa Cruz Biotech. 1:300); and Mouse anti-CD44 (ab6124, abcam, 1:250). Primary antibodies were incubated for 4 days at 4°C, followed by application of secondary antibodies (anti-mouse 647, Cat#A32787, RRID: AB_2762830, Invitrogen; anti-rat 594, A21209, RRID: AB_2535795, Invitrogen) and another 4-day incubation at 4°C. Finally, 4’,6-diamidino-2-phenylindole (DAPI, D9542, Sigma-Aldrich) was added and stained for 1 day. Samples were then submerged in an optical clearing solution overnight and embedded in a 1% agarose gel for imaging with a Leica TCS SP8 confocal microscope.

### Single-cell RNA sequencing

Cyborg and control SC-islets were dissociated into single cells as described previously^7,8,30^. Then, the single cells were suspended in DPBS (without Ca^2+^ and Mg^2+^) with 0.04% bovine serum albumin (Sigma) at a concentration of 1000 cells per microliter. Library preparation and sequencing were carried out at the Bauer Sequencing Core facility at Harvard University. The 10X Genomics Chromium Single Cell 3′ v3 Reagent Kit was employed to prepare samples following the experimental protocol, as guided by the 10X Genomics Single Cell Protocols Cell Preparation Guide. Subsequently, the prepared samples underwent sequencing on the Illumina NovaSeq platform, with the following sequencing specifications: platform - NovaSeq S4 full flow cell, read length - 50, and read type - paired end.

### Single-cell RNA-seq data analysis

The read alignments were first performed with Cell Ranger (10x Genomics) to the reference human genome GRCh38. Then the R package Seurat was used to analyze the scRNA-seq data. The cells were first filtered for quality control (mitochondrial reads <20%, genes detected >800 and < 12500). Then the cell gene expression matrices were normalized and scaled using the “NormalizedData()”, “FindVariableFeatures()” and “ScaleData()” functions. Next, dimensional and clustering analysis was performed with “RunPCA()”, “FindNeighbors()” and “FindClusters()” functions, followed by “RunUMAP()” function to visualize the data. To annotate cell-types, we compared marker genes for each cluster with those of previous publications^7,14^.

For functional enrichment and protein-protein interaction analysis before and after circadian entrainment experiment, the differentially expressed genes for each cell-type (e.g., SC-α, β, and EC) were used as input for the Metascape software (https://metascape.org/gp/index.html#/main/step1). The graph visualizations were then performed with the Cytoscape software^53^.

For pseudotime analysis, gene expression from each cell subtype was first subsetted and then the PHATE (https://github.com/KrishnaswamyLab/phateR) algorithm was performed to project the gene expression into 2D PHATE space using the “phate()” function. The Slingshot (https://github.com/kstreet13/slingshot) algorithm was then performed to infer the pseudotime trajectory with “slingshot()”.

### Spike sorting

Electrical recordings processing and spike sorting were performed using MountainSort^31^ and SpikeInterface (version 0.98.2)^32^. The raw electrical recordings underwent bandpass filtering within the frequency range of 300-3,000 Hz. Common average reference was applied to reduce the common-mode noise from the recording. Each recording session was processed individually for spike sorting. Additionally, recordings from each of the 64 channels was sorted separately. To extract spike waveforms, a spike detection threshold was applied, set at 5.5 times the standard deviation away from the mean. The output of spike sorting was manually curated to remove noisy units, which were excluded if the raw electrical recording trace is noisy, or if the number of waveforms contained within the unit is sufficiently small (e.g., once per recording trace).

### Cell type annotation

Averaged electrical firing rates of spike-sorted units were computed for each minute recorded under 2.8 mM and 20 mM glucose. Two-tailed, unpaired Welch’s t-tests were then conducted on the per-minute firing rates under the two glucose conditions for each unit. A unit was annotated as an SC-α cell if it exhibits higher firing rates in 2.8 mM glucose with a p-value < 0.05, or as a SC-β cell if it exhibits higher firing rates in 20 mM glucose with a p-value < 0.05. If a unit did not exhibit significantly different firing rates between 2.8 mM and 20 mM glucose, it was labeled as “other”. To further classify an SC-α cell into α1 or α2 states, we computed the ratio of its averaged per-minute firing rate in 2.8 mM glucose over its rate in 20 mM glucose. Then the multiple of the median (MoM) was computed as a measure of how far an individual SC-α cell firing rate ratio deviated from the median across the population. SC-α cells with MoM > 2 were assigned to an α1 state and those with MoM < 2 to an α2 state. To further classify an SC-β cell into β1 or β2 states, the ratio of its averaged firing rate in 20 mM glucose over that in 2.8 mM glucose was computed. Then SC-β cells with MoM > 2 were assigned to a β1 state and those with MoM < 2 to a β2 state. A small firing rate offset was added to the denominator when calculating firing rate ratios to avoid zero-division errors. All computations were carried out using Python 3.8 and Scipy 1.10.1^54^.

### Rhythmicity analysis

Rhythmicity of hormone level and voltage spike firing rate time series measurements was evaluated with the RAIN R package^55^, which uses non-parametric Mann-Whitney U tests to compare the ranks of measured values against those of alternative waveforms without assumptions of waveform shape or symmetry, and calculates a Benjamini-Hochberg corrected p value. We implemented RAIN with default parameters and “period = 24, deltat = 4”. We specified “measure.sequence” for measurements with varying number of replicates and “na.rm = TRUE” for those with NAs. We used “method = independent” for electrical recording and “method = longitudinal” for hormone level measurements, representing a time series sampled from the same bulk cell culture. Sinusoidal regressions of detrended hormone and electrical measurements were plotted for visualization purposes. Comparable trends and similar statistical significance estimates were obtained using harmonic regression with the Harmonic Regression R library^56^ or using the eJTK Rhythmicity test^57^. For plotting, time series measurements were detrended by subtracting a linear regression fit, considering both measurement means and SEMs to normalize whitin the same scale. Hormon level measurements were computed using BioDare2^58^ and electrical recording processing was done using Python 3.8 and Scipy 1.10.1.

### Waveform analysis

Waveform analysis was performed using Scanpy 1.9.5^57^ and PHATE^41^. Spike waveforms of all concerned units were first aligned by inverting waveforms with troughs preceding peaks. All waveforms from the same measured timepoint were averaged to obtain template waveforms for the 18 measurements over 72 hours. Templates in between two timepoints were linearly interpolated. Then the original waveforms, averaged template waveforms, and interpolated waveforms were jointly projected onto the same 3D PHATE space.

## Data availability

The scRNA-seq data are available in the Single Cell Portal. The electrical data are available in the GitHub repository.

## Acknowledgements

We thank Douglas A. Melton for his generous support during the initial stage of this project. We acknowledge the support from NIH/NIDDK 1DP1DK130673, the Human Islet Research Network (U24DK104162) and from a pilot award from the Diabetes Research Center at the University of Pennsylvania (P30DK19525) to J.R.A-D., who was also supported by grants to Douglas A. Melton from the JDRF (5-COE-2020-967-M-N), and the JPB Foundation (award no. 1094).

## Author contributions

J.L., J.RA.-D., Q.L., and R.L. conceived of the experimental design. R.L. designed, fabricated, and characterized the properties of the devices. J.RA.-D., S.D.P., and I.G.S. performed SC-islets differentiation and ELISA experiments. Q.L. performed cell cultures, device integrations, immunofluorescence staining experiments, and prepared samples for scRNA-seq. Q.L. and R.L. performed circadian entrainment experiments and electrophysiological recordings. Z.L. analyzed scRNA-seq data. X.Z., Q.L., and I.G.S. analyzed electrophysiological recordings data. Q.L., Z.L., J.RA.-D., and J.L. wrote the manuscript. All authors have reviewed the final version of the manuscript. J.L. and J.RA.-D. supervised the study.

## Additional information

Correspondence and requests for materials should be addressed to J.L. and J.RA.-D.

---

**Extended Data Fig. 1.**
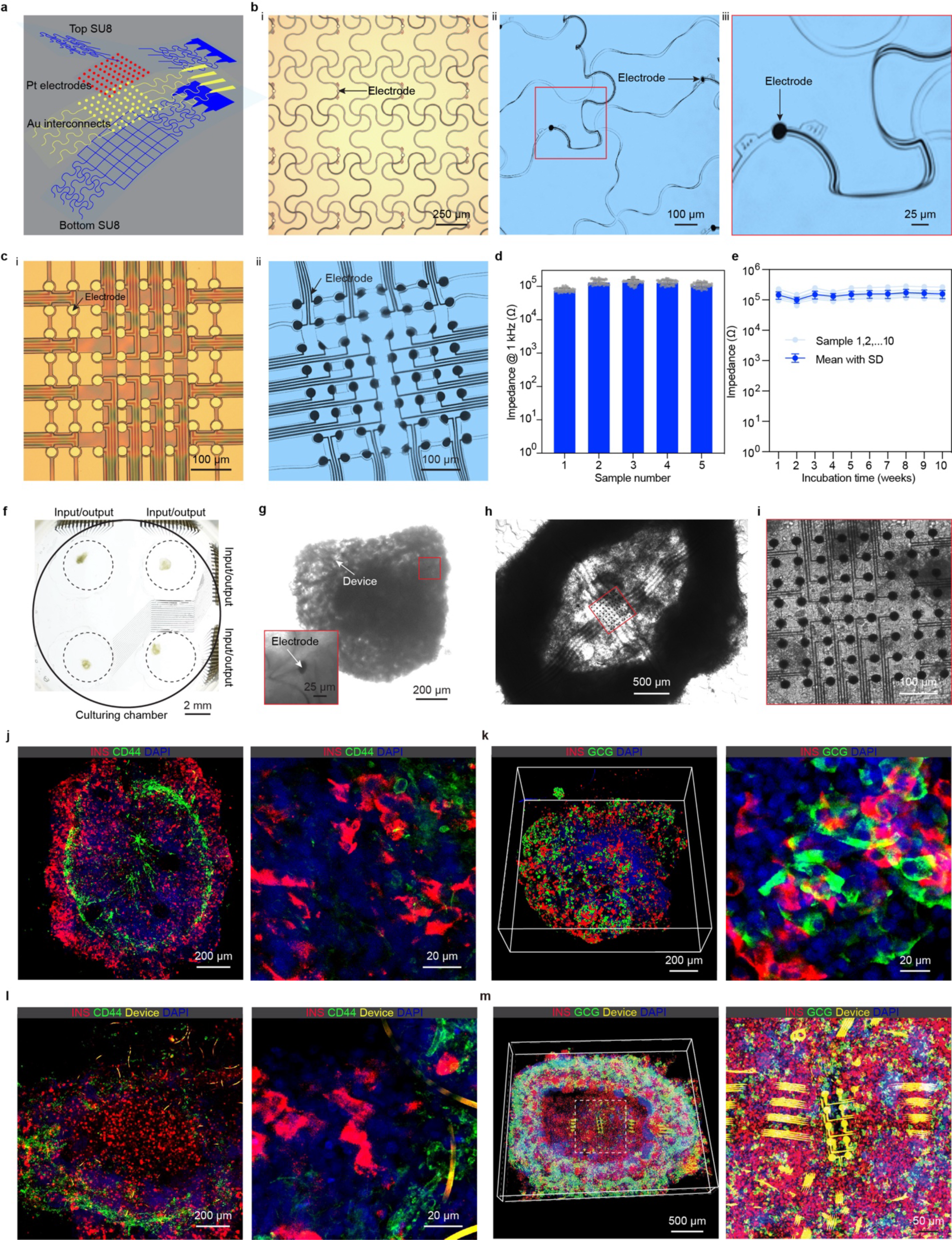
Design, fabrication, and integration of stretchable mesh electronics into cyborg islets. **a**, Schematic illustrating the multilayer structure of stretchable mesh electronics. **b**, Representative bright-field (BF) images of unreleased (**i**) and released (**ii**) 16-channel stretchable mesh electrodes. Zoom-in image (**iii**) shows the individual electrode, fluorescence bar code, and twisted stretchable interconnects. **c**, Representative BF images of representative unreleased (**i**) and released (**ii**) 64-channel high-density electrode arrays in stretchable mesh electronics. **d**, Average electrochemical impedance of electrodes at 1 kHz (n = 64 electrodes for each sample device, values are mean ± SEM). **e,** Average impedance of electrodes at 1 kHz as a function of incubation time in PBS at 37°C (n = 10 samples, values are mean ± SD). **f**, Optical photograph of a representative culturing chamber with four cyborg islets. **g**, BF phase images show a representative cyborg islet integrated with 16-channel stretchable mesh nanoelectronics. Inset shows the zoom-in view of red box-highlighted region showing an electrode embedded in the islet. **h-i**, BF phase image of a representative cyborg islet integrated with 64-channel stretchable mesh electronics (**h**) and zoom-in view (**i**) show the embedded high-density electrode array. **j-k**, Fluorescence images of cleared, immunostained control SC-islets (without device integration) with 16-channel stretchable mesh nanoelectronics. Zoom-in views show cell morphologies and cell-device coupling, respectively. Insulin (INS, red), glucagon (GCG, green), CD44 (green), device (yellow), and DAPI (blue). **l-m**, Fluorescence images of cleared, immunostained cyborg SC-islets with 16-channel stretchable mesh nanoelectronics (**l**) and 64-channel stretchable mesh nanoelectronics (**m**). Zoom-in views show cell morphologies and cell-device coupling, respectively. Insulin (INS, red), CD44 (green), glucagon (GCG, green), device (yellow), and DAPI (blue).

**Extended Data Fig. 2.**
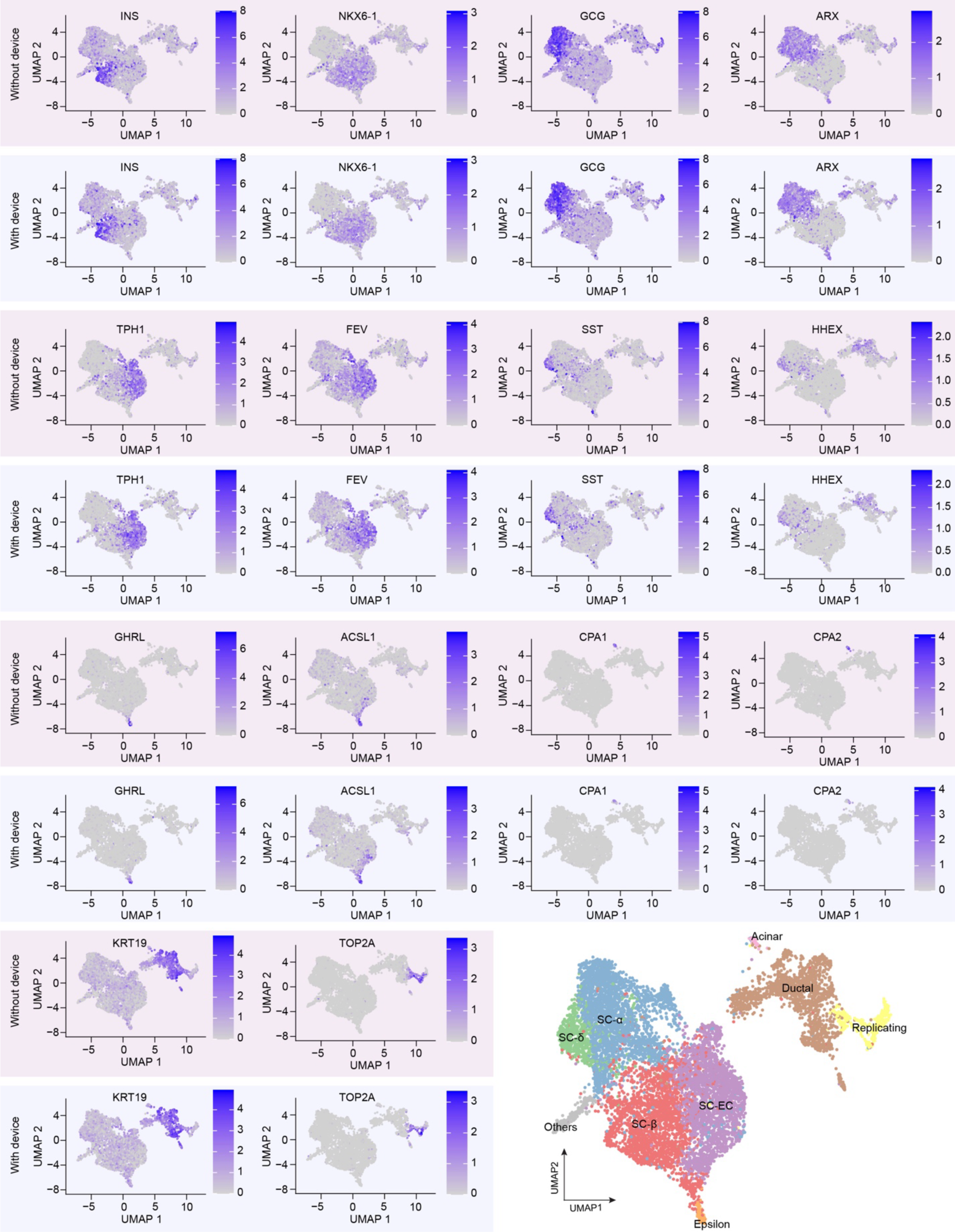
Expression of gene markers in cyborg and control islets. 2D UMAP visualization of clustering of single-cell RNA expression profiles for the indicated marker genes in cyborg (with device) and control (without device) islets. Cells are colored by gene expression level.

**Extended Data Fig. 3.**
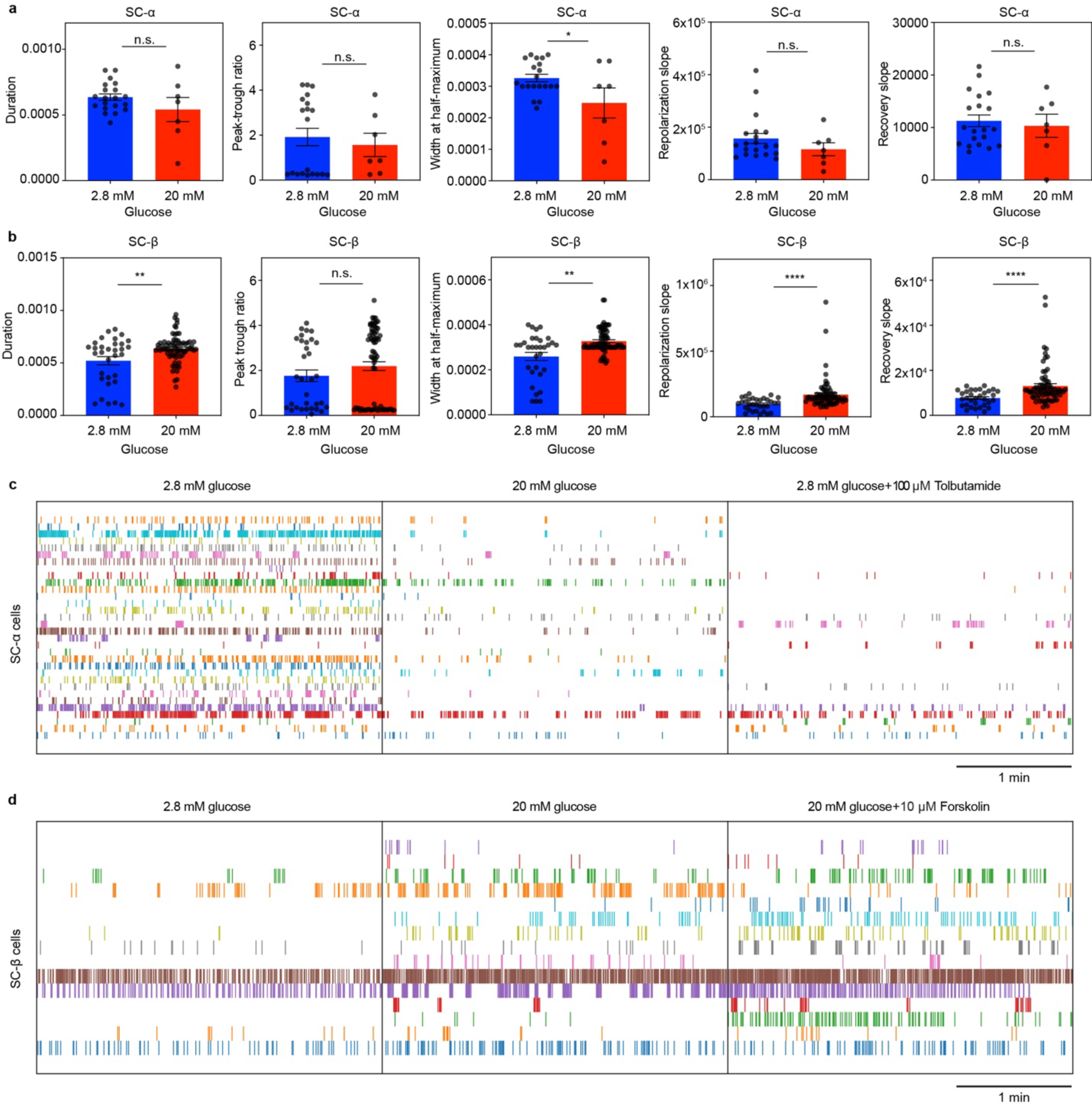
Analysis of cell type-specific electrical features from cyborg islet recordings. **a-b**, Electrical features from SC-α (**a**) and SC-β (**b**) cell recordings. Spike duration, peak-trough ratio, width at half-maximum, repolarization slope, and recovery slope were analyzed and compared between 2.8 mM and 20 mM glucose incubations. n.s., not significant; * p < 0.05; ** p < 0.01; *** p < 0.001; **** p < 0.0001 from two-tailed, unpaired t test. **c,** SC-α cell spike trains recorded at 2.8 mM glucose, 20 mM glucose, and 2.8 mM glucose + 100 μM tolbutamide (N = 32 cells pooled from n = 3 cyborg islets). **d**, SC-β cell spike trains recorded at 2.8 mM glucose, 20 mM glucose, and 20 mM glucose + 10 μM forskolin (N = 15 cells pooled from n = 3 cyborg islets).

**Extended Data Fig. 4.**
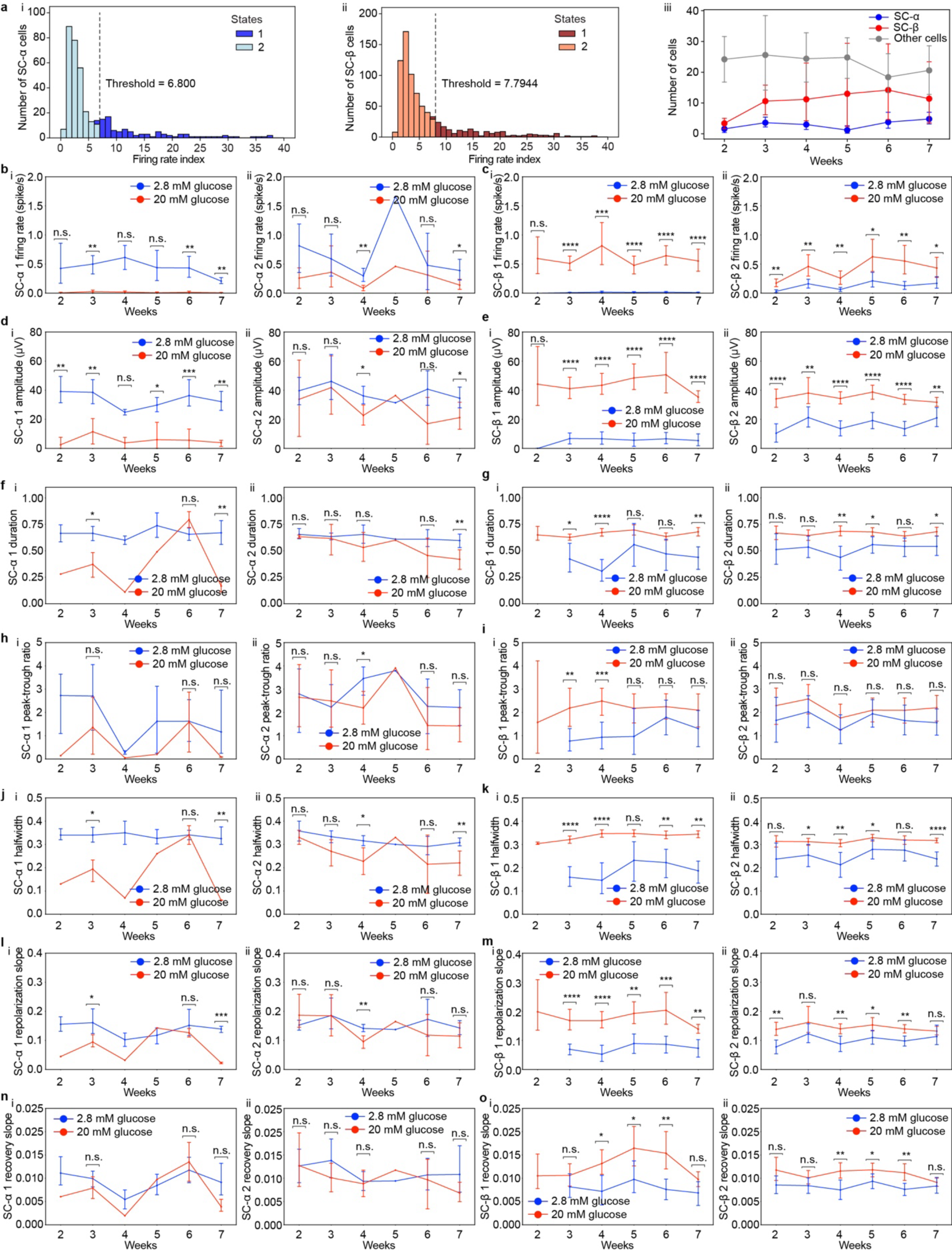
Analysis of cell type-specific electrical features from cyborg islet long-term recordings. **a,** The multiple of the median (MoM) as a measure of how far an individual SC-α (i) or -β (ii) cell firing rate ratio deviated from the median across the population for further classifying cell state 1 and 2 (See Methods). The number of electrically recorded SC-α, SC-β and other cells from week 2 to week 7 after device integration (iii). Data are mean ± SEM of N = 380 SC-α, 800 SC-β, and 2776 other cells pooled from n = 6 cyborg islets. **b,** Spike firing rates for SC-α 1 (i) and -α 2 (ii) cells at 2.8 mM and 20 mM glucose from week 2 to week 7 after device integration. **c,** Spike firing rates for SC-β 1 (i) and - β 2 (ii) cells at 2.8 mM and 20 mM glucose from week 2 to week 7 after device integration. **d,** Amplitudes for SC-α 1 (i) and -α 2 (ii) cells at 2.8 mM and 20 mM glucose from week 2 to week 7 after device integration. **e,** Amplitudes for SC-β 1 (i) and -β 2 (ii) cells at 2.8 mM and 20 mM glucose from week 2 to week 7 after device integration. **f-o**, Electrical features including spike duration (**f,g**), peak-trough ratio (**h,i**), width at half-maximum (**j,k**), repolarization slope (**l,m**), and recovery slope (**n,o**) for SC-α 1 & β 1 (i) and SC-α 2 & β 2 (ii) cells were analyzed. Plots show mean ± SEM of N = 33 SC-α 1 and 57 SC-α 2, 126 SC-β 1 and 193 SC-β 2 cells pooled from n = 5 cyborg islets. Two-tailed, unpaired Welch’s t-tests were conducted under the two glucose conditions.

**Extended Data Fig. 5.**
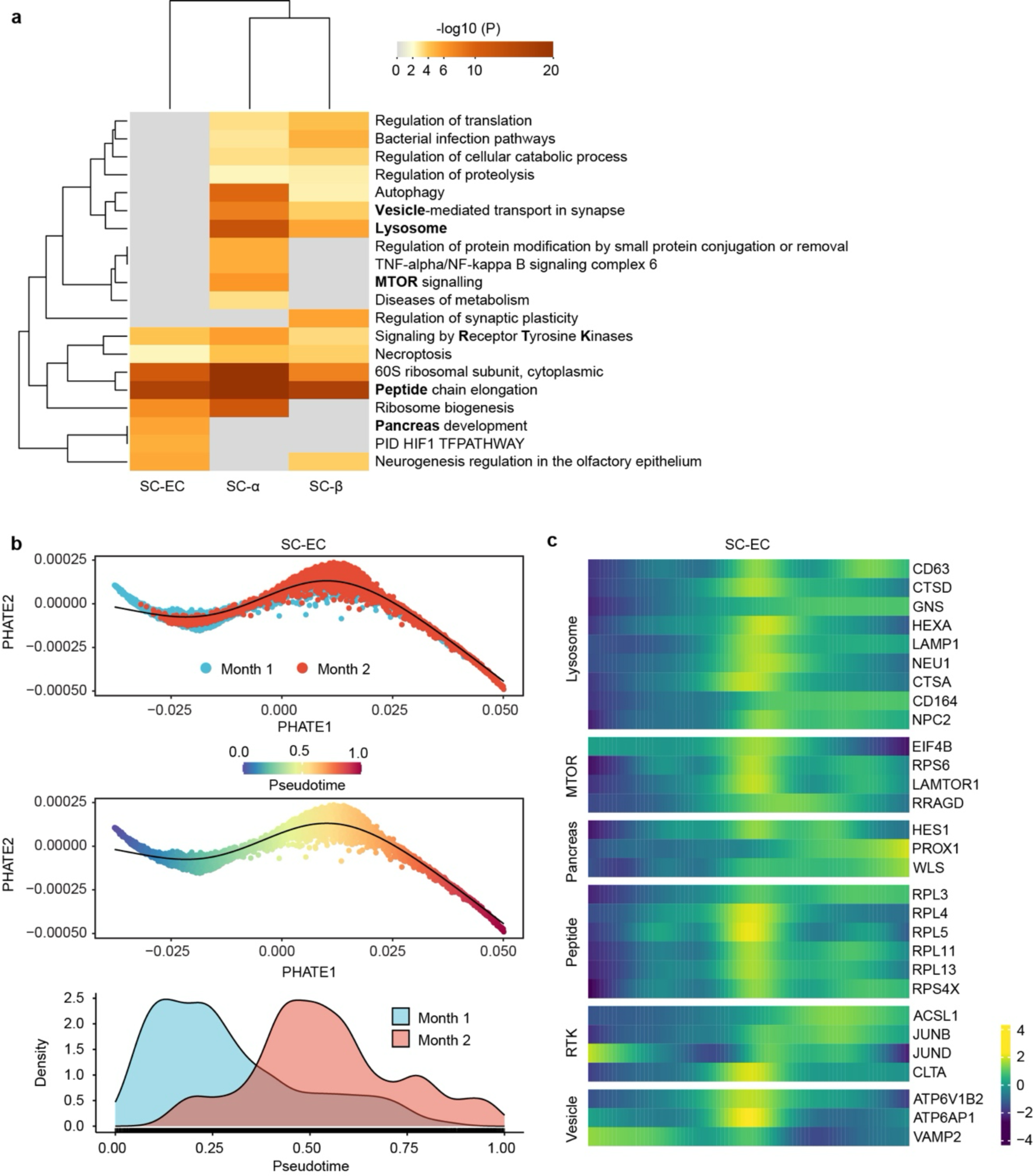
Early cyborg islet maturation characterized by sc-RNA seq. **a**, Heatmap showing biological functions enriched among genes upregulated in SC-α, SC-β, or SC-EC cells in later (month 2) versus earlier (month 1) pseudotime. **b**, Early maturation trajectory of SC-EC cells using 2D PHATE visualization, by month (top) or inferred pseudotime ordering (middle), and pseudotime distributions of month 1 and month 2 islet cells (bottom). **c**, SC-EC pseudotime-dependent gene expression programs along with their enriched pathways, which are also highlighted in bold in panel (**a**). Heatmap shows the row z-scored expression of selected dynamic genes along SC-EC pseudotime.

**Extended Data Fig. 6.**
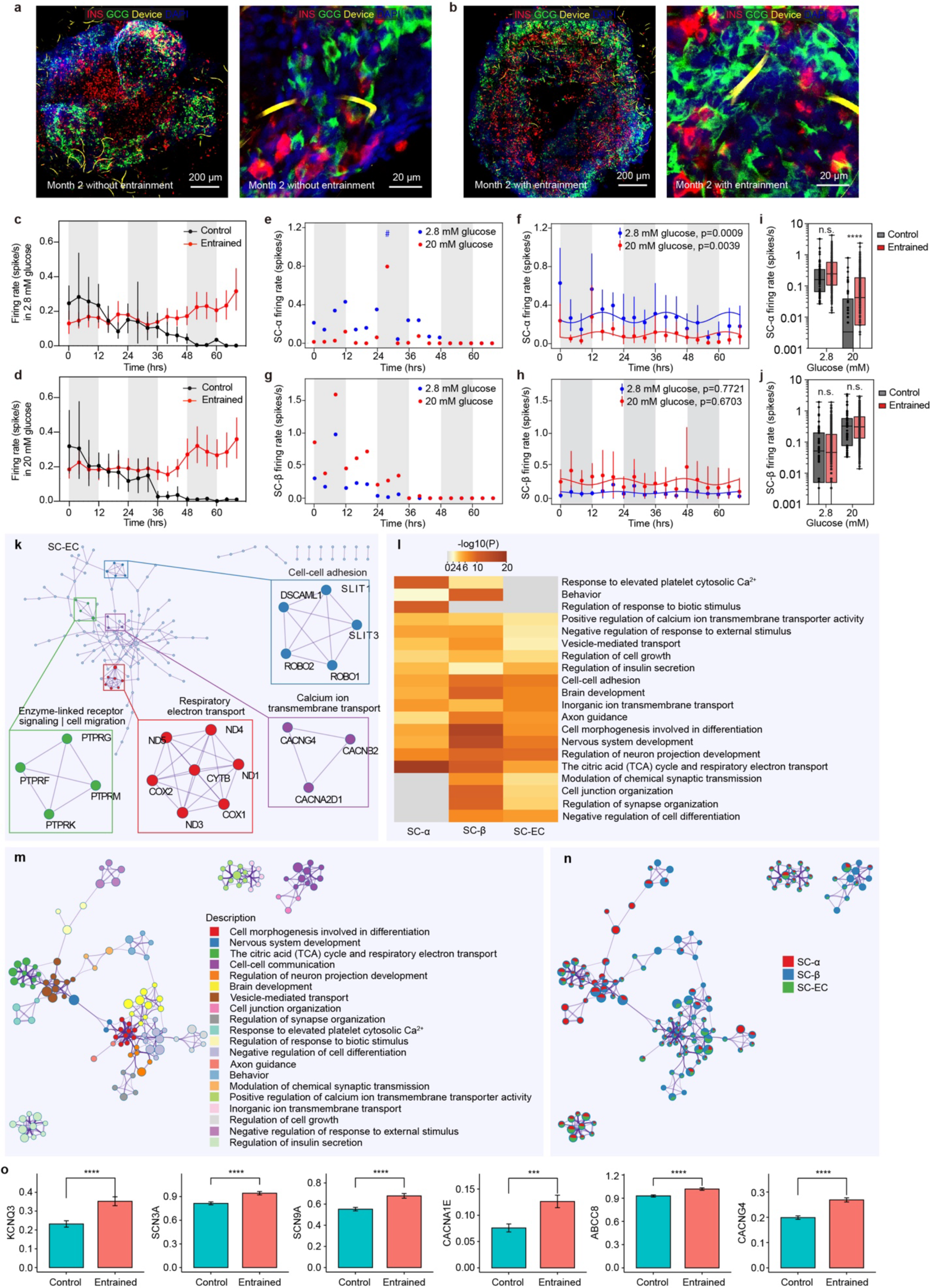
SC-α and SC-β cell maturation triggered by circadian entrainment. **a-b**, Fluorescence images of tissue cleared, immunostained cyborg islets without (**a**) and with (**b**) circadian entrainment at month 2 after device integration. Zoom-in views show cell morphologies and cell-device coupling. Insulin (INS, red), Glucagon (GCG, green), device (yellow), and DAPI (blue). **c-d**, Spike firing rates at 2.8 mM (**c**) and 20 mM (**d**) glucose recorded over 72 hours after cyborg islet diurnal feeding (entrained) or mock (control) entrainment. Data are mean ± SEM across N = 220 recording channels pooled from 5 independent entrained cyborg islets and across N = 50 recording channels from a control islet. **e-h**, Spike firing rate rhythms of SC-α cells during mock (control) entrainment (**e**) and after diurnal feeding (entrained) (**f**), and SC-β cells during mock (control) entrainment (**g**) and after diurnal feeding (entrained) (**h**) at 2.8 mM and 20 mM glucose recorded over 72 hours. P-values, RAIN rhythmicity test^55^ (See Methods). **i-j**, Spike firing rates of SC-α (**i**) and SC-β (**j**) cells at 2.8 mM and 20 mM glucose after cyborg islet diurnal feeding (entrained) or mock (control) entrainment. Data are mean ± SEM across N = 200 recording channels over 72 hours pooled from 5 independent entrained cyborg islets and across N = 50 recording channels over 72 hours from a control islet. **k**, Biological functions enriched among protein-protein interaction complexes identified by the MCODE^60^ algorithm based on genes upregulated in SC-EC cells in circadian-entrained versus control cyborg islets. The top 4 interaction complexes identified (calcium ion transmembrane transport, respiratory electron transport, cell-cell adhesion, and enzyme-linked receptor signaling | cell migration) are highlighted. **l**, Heatmap showing biological functions enriched among genes upregulated in SC-α, SC-β, or SC-EC cells in circadian-entrained versus control cyborg islets. **m-n**, Network visualization of enriched biological pathways among genes upregulated in circadian-entrained versus control cyborg islets, colored by enrichment cluster terms (**m**) or by cell type (**n**). **o**, Barplots showing expression of selected genes in control versus circadian-entrained cyborg islets. Data are mean ± s.e.m., *** p<0.001, **** p<0.0001.

